# dubTAGs enable on-demand stabilization for tunable and reversible control of endogenous protein levels

**DOI:** 10.64898/2026.09.11.751045

**Authors:** Sunil Guharajan, Xiangyang Song, Sachi Sengupta, Qiong Wu, Wenyi Wei, Yan Xiong, Jian Jin, Sahin Naqvi

**Affiliations:** Division of Gastroenterology, Hepatology and Nutrition, Department of Pediatrics, Boston Children’s Hospital and Harvard Medical School, Boston, MA, USA; Broad Institute of Massachusetts Institute of Technology and Harvard, Cambridge, MA, USA; Mount Sinai Center for Therapeutics Discovery, Departments of Pharmacological Science, Oncological Science, and Neuroscience, The Mount Sinai Tisch Cancer Center, Icahn School of Medicine at Mount Sinai, New York, NY, USA; Department of Pathology, Beth Israel Deaconess Medical Center, Harvard Medical School, Boston, MA, USA

## Abstract

Precise and rapid control over cellular protein levels is essential to dissect complex biological systems. Chemical genetic approaches such as dTAG, in which a target is fused to a degron tag (FKBP12^F36V^) and degraded upon small molecule-mediated recruitment of E3 ligases, have enabled rapid and tunable control over protein abundance. However, no analogous tool exists to precisely increase protein levels and actively reverse dTAG-mediated degradation. Here, we developed heterobifunctional small molecules (dubTAGs) that stabilize FKBP12^F36V^-tagged proteins by recruiting endogenous deubiquitinases. Utilizing stem cell-derived cranial neural crest cells (CNCCs) in which the transcription factors SOX9 or TWIST1 are endogenously tagged with FKBP12^F36V^, we identified OTUB1- or USP7-recruiting heterobifunctional molecules that demonstrated effective target stabilization and ternary complex formation. We demonstrate that dubTAG-mediated protein stabilization is dependent on deubiquitinase recruitment, target-specific, and can tunably and rapidly reverse dTAG-mediated degradation. We applied dubTAGs to assess how stabilizing endogenous SOX9 impacts chromatin accessibility in CNCCs, finding both monotonic and non-monotonic regulatory element responses that are driven by distinct sequence features. dubTAGs are readily applicable tools for investigating the effects of elevated protein levels and tunably reversing targeted degradation, enabling new approaches to study protein dosage effects in development, disease, and therapeutic discovery.

## Introduction

Precise control over both the levels and temporal dynamics of cellular proteins is essential for faithful development and homeostasis. Imbalances in protein stoichiometry, also termed dosage sensitivity, is a frequent cause of disease^1^. Haploinsufficiency, caused by loss of one functional allele and a ∼50% reduction in protein levels, is a frequent cause of developmental disorders. Similarly quantitative increases in protein levels can also be detrimental: single-gene duplications or aneuploidies that result in ∼50% increases in protein levels have been associated with developmental disorders and cancer outcomes, a phenomenon termed triplosensitivity^2,3^. Non-monotonic gene dosage effects, where both deletions and duplications have significant effects in the same direction, have also been observed for human complex traits^4^. Beyond steady-state abundance, temporal dynamics of protein levels are well-established determinants of cellular behavior, with the timing and duration of activity shaping outcomes during processes such as signaling and cell-fate specification^5^.

Dissecting the mechanisms that underlie sensitivity to both decreases and increases in protein dosage requires tools that can rapidly, reversibly, and bidirectionally tune target protein levels. Targeted protein degradation, in which small molecules induce selective degradation of a protein of interest (POI) by hijacking cellular ubiquitin-proteasome signaling, has emerged as a highly promising approach towards this end^6 7^. Although a range of targeted protein degradation technologies have been developed, the degradation TAG (dTAG) system has been particularly widely adopted in mammalian settings. In this system, the POI is fused to the degron tag FKBP12^F36V^ and degraded by the addition of heterobifunctional small molecules that bind FKBP12^F36V^ and recruit endogenous E3 ligases^8,9^. Indeed, since its inception in 2018, at least 72 studies have utilized dTAG in both *in vitro* and *in vivo* settings, and the rate of such publications is increasing over time (Figure 1, Table S1). Despite the success of dTAG, it is fundamentally based on reducing protein levels, and can therefore only address the effect of decreased protein levels. While reversibility of dTAG has been demonstrated in some settings through washout experiments, the degree of this reversibility is not controllable and its kinetics are dependent on cellular dilution from division and the off-rate of the dTAG molecule from FKBP12^F36V^. The Shield system has been developed for selective stabilization and fast induction of target proteins by endogenous tagging with a FKBP12-derived destabilization domain that is stabilized upon small molecule binding^10^. However, the Shield approach requires the default state of the tagged protein to be degraded, making it challenging to apply to essential proteins required for cellular fitness and cell-fate transitions. Overcoming these limitations would enable studies of how rapid restoration of endogenous proteins affects cellular processes.

**Figure 1.**
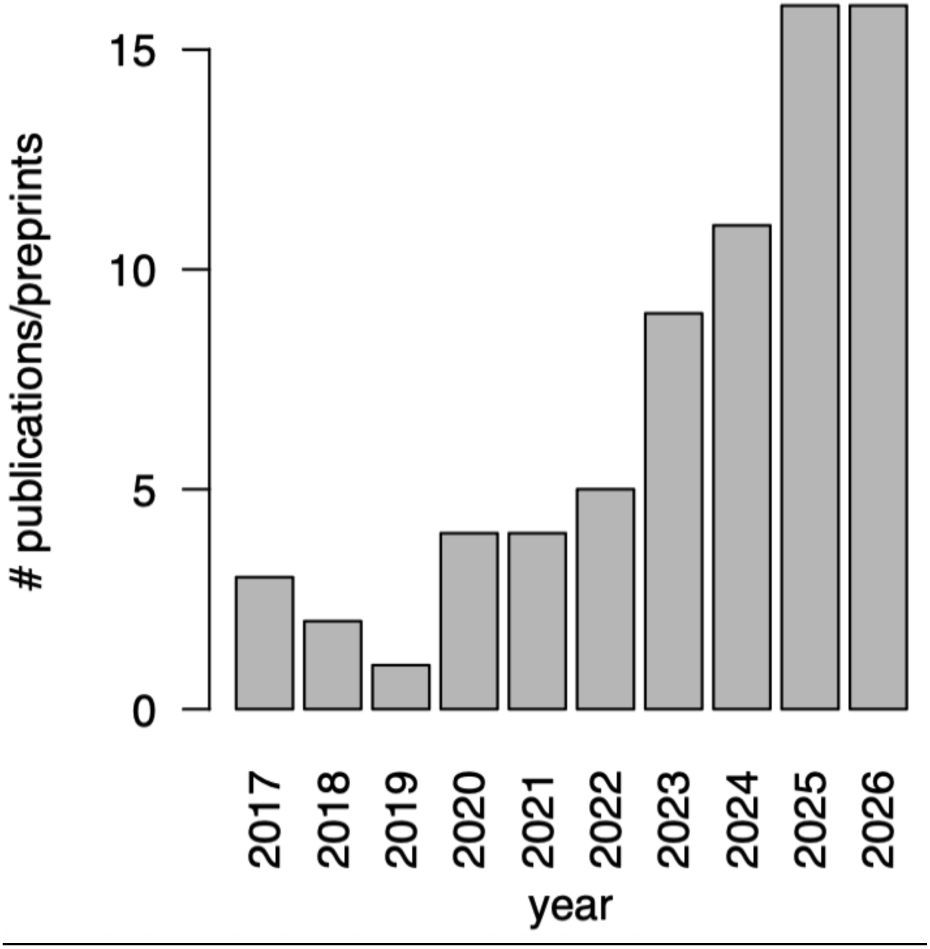
Number of publications or preprints utilizing the dTAG system by year. 2026 is a partial year.

We sought to develop a tool that could specifically increase endogenous protein levels and tunably reverse targeted protein degradation by recruiting endogenous deubiquitinases. Prior studies by Henning and colleagues pioneered the concept of targeted protein stabilization to reverse disease-causing aberrant ubiquitination^11^, but such molecules are target-specific and therefore not general tools for probing the effects of protein stabilization and resulting dosage increases or reversal of degradation. We reasoned that heterobifunctional small molecules that recruit endogenous deubiquitinases to FKBP12^F36V^-tagged POIs would increase stability and lead to higher levels. Such a system would also allow for the active reversal of dTAG-mediated degradation and provide a broadly applicable tool that would unlock targeted stabilization for the many existing cellular reagents in which a variety of POIs have been tagged with FKBP12^F36V^.

## Results

### Screening heterobifunctional molecules in FKBP12^F36V^-tagged cells reveals candidates that increase intracellular protein concentrations

We first sought to identify a system that could assess the ability of heterobifunctional small molecules (dubTAGs) that recruit deubiquitinases to FKBP12^F36V^-tagged, endogenous proteins (Figure 2A). We previously generated human embryonic stem cells (hESCs) in which both alleles of the transcription factor SOX9 were tagged with FKBP12^F36V^ and the fluorescent protein mNeonGreen, allowing for quantitative assessment of SOX9 protein levels^12^. These hESCs can be differentiated into cranial neural crest cells (CNCCs), the main progenitors of the face, where SOX9 is highly expressed and is a major determinant of chromatin accessibility and gene expression. Previous studies have reported the half-life of SOX9 protein to be between 3 and 7 hours depending on cellular context^13,14^; we therefore reasoned that increasing SOX9 half-life by treating SOX9-tagged cells with dubTAGs for 24-48h should result in discernable increases in steady state protein levels.

**Figure 2.**
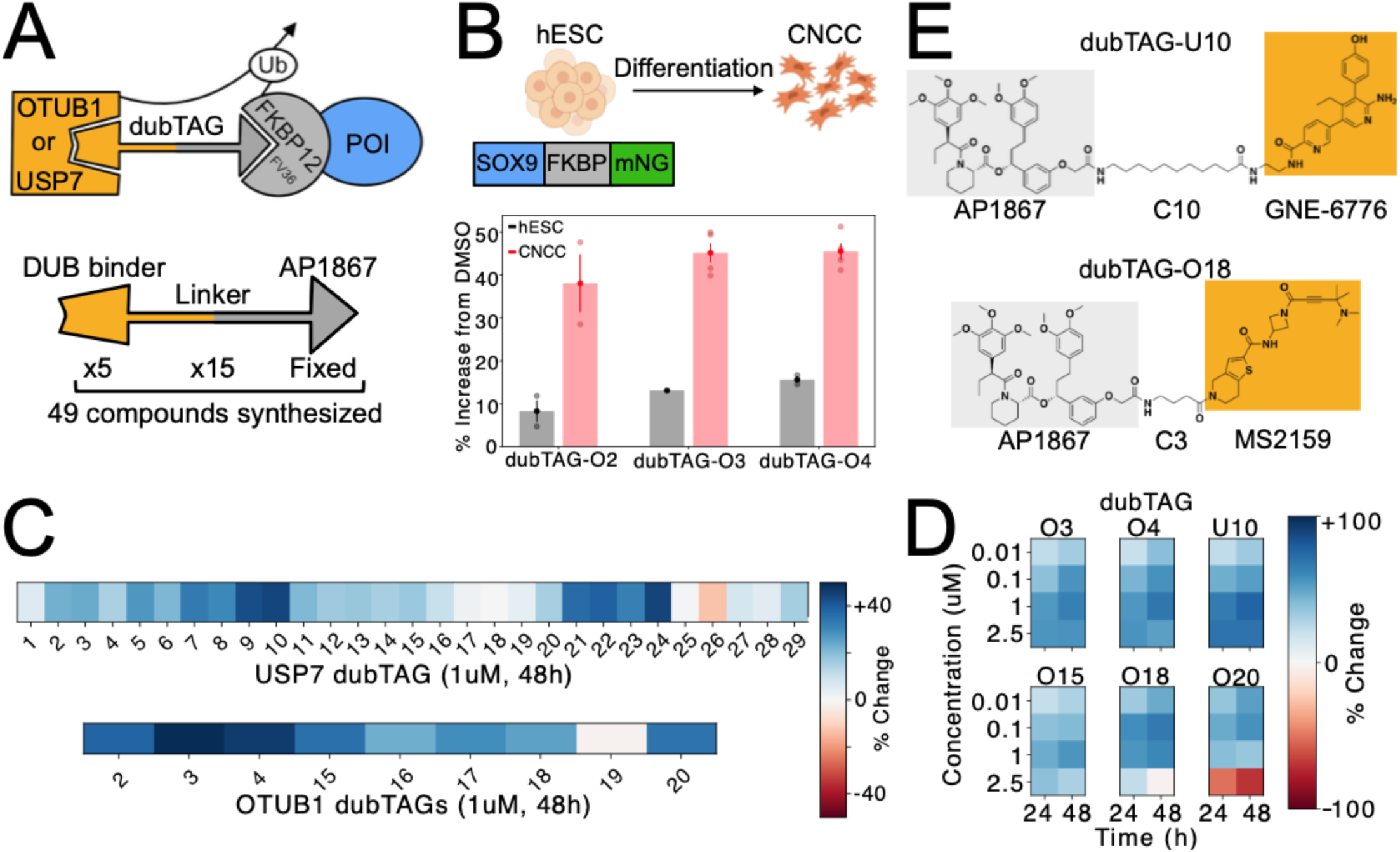
Identification of heterobifunctional small molecules that mediate targeted stabilization of FKBP12^F36V^-tagged proteins (dubTAGs). (A) Top, strategy for regulating stabilization and levels of proteins of interest (POI) by recruitment of E3 ligases and cellular deubiquitinases (DUB) OTUB1 or USP7. Bottom, schematic of dubTAG compounds synthesized. (B) Effects of selected dubTAGs on SOX9 levels in human embryonic stem cells (hESCs) or hESC-derived cranial neural crest cells (CNCCs) when applied at 1 μM for 48h. Light points represent independent biological replicates, dark point and error bars indicate mean and standard error. (C) Effects of all screened dubTAGs in CNCCs on SOX9 levels. Data represent the mean effect size with minimum n=2 replicates per compound. (D) Effects of selected dubTAGs on SOX9 levels in CNCCs with multiple concentrations (y-axis) and treatment times (x-axis). Data represents the mean effect size of minimum n=2 replicates per compound. (E) Chemical structures of top candidates after screening. dubTAG-U10 utilizes the USP7 binder GNE-6776 while dubTAG-O18 uses a novel OTUB1 binder MS2159.

We first designed and synthesized a series of putative dubTAGs, each containing three parts: a flexible linker region with variable carbon-or polyethylene glycol (PEG) lengths connecting the FKBP12^F36V^ binder AP1867^8^ to MS5105, the small-molecule binder for the deubiquitinase enzyme OTUB1 we developed previously^15^ (Table S2). Next, these initial 14 compounds (O1 – O14) were screened in SOX9-tagged hESCs with a fixed concentration and duration (1 μM for 48h) and assessed resulting SOX9 levels by flow cytometry for mNeonGreen (GFP). We observed a 12-16% increase in SOX9 levels among the top performing candidates (O2 – O4) (Figure S1). We reasoned that the relatively modest dubTAG effect sizes observed could be due to the fact that SOX9 is lowly expressed in hESCs, resulting in a slow baseline ubiquitination and degradation rate (which in turn sets the maximum dubTAG-mediated increase in steady state levels). We therefore used a robust protocol^16^ to differentiate the SOX9-tagged hESCs into CNCCs and profiled a subset of dubTAGs at the same concentration and treatment duration (Figure 2B). We found larger effect sizes (40-45% increase) of the same dubTAGs in CNCCs, likely reflecting the larger contribution of ubiquitin-mediated proteasomal degradation to SOX9 steady state levels.

Having established SOX9-tagged CNCCs as an optimal screening context, we synthesized two additional sets of OTUB1-recruiting dubTAGs (Table S2). These compounds were derived from the two next-generation OTUB1 binders recently developed in our lab: MS8572^17^ for O15 – O17 and MS2159^18^ for O18 – O20. Each OTUB1 binder was conjugated to AP1867 using three-to five-carbon linkers, similar in length to those used in the lead compounds. We also synthesized two series of USP7 recruiting dubTAGs (U1 – U29) using the previous reported USP7 ligands GNE-6776^19^ and USP7i-#4a^20^ (Table S2), both of which have previously been employed for DUBTAC development^21^. We next screened these novel 35 USP7 and OTUB1-recruiting dubTAGs as well as the initial 3 hits (O2 –O4) at a concentration of 1 μM for 48h. We observed several candidates that achieved a 40-50% increase in SOX9 levels (Figure 2C); From these compounds, we selected one GNE-6776-derived USP7-recruiting dubTAG (U10), two MS5105-derived OTUB1-recruiting dubTAGs (O3 and O4), one MS8572-derived OTUB1-recruiting dubTAG (O15), and two MS2159-derived OTUB1-recruiting dubTAGs (O18 and O20) for further characterization across four concentrations and two treatment durations (24 and 48h). Most dubTAGs showed some effect by 24h but increased in effect size by 48h, consistent with a mechanism of SOX9 protein accumulation from increased half-life (Figure 2D). Furthermore, all six dubTAGs showed reduced efficacy at 2.5 μM, demonstrating a hook effect characteristic of heterobifunctional molecules^22^. From these data, we identified two top candidates: 1) dubTAG-U10 (MS142) with maximal effect at 1 μM, and 2) dubTAG-O18 (MS225), which achieved maximal effect at 0.1 μM for 48h treatment.

To assess whether dubTAGs could stabilize target proteins beyond SOX9, we applied dubTAG-O18 and dubTAG-U10 to CNCCs in which the TF TWIST1 has been tagged with FKBP12^F36V^ and the V5 epitope^23,24^. Treatment with dubTAG-O18 consistently resulted in a ∼30% increase in TWIST1 levels as measured by intracellular flow cytometry, while the effect of dubTAG-U10 was overall minor and varied by differentiation batch (Figure S2). Together, these data demonstrate effective stabilization of FKBP12^F36V^-tagged, endogenous proteins through a deubiquitinase recruitment strategy via dubTAGs. Our results further suggest that the most effective dubTAG (or recruited deubiquitinase) may vary based on the target protein, underscoring the importance of a strategy that specifically recruits individual deubiquitinases.

### dubTAGs induce ternary complex formation

We next assessed whether dubTAGs induce ternary complex formation between FKBP12^F36V^ and deubiquitinases by applying the NanoLuc Binary Technology (NanoBiT) complementation system^25^, in which proximity of the SmBiT and LgBiT peptides induces luciferase signal (Figure 3A). Focusing on dubTAG-O18, we expressed OTUB1-SmBiT and LgBiT-FKBP12^F36V^ in HEK293 cells under the control of the CMV promoter and assessed the increase in luciferase signal upon 24h treatment with either dubTAG-O18 or equimolar binders (MS2159 and AP1867) compared to DMSO. We observed small increases (less than 1.5-fold) with CMV-expressed OTUB1, although dubTAG-O18 showed consistently higher fold-changes than equimolar binders across doses (Figure 3B inset).

**Figure 3.**
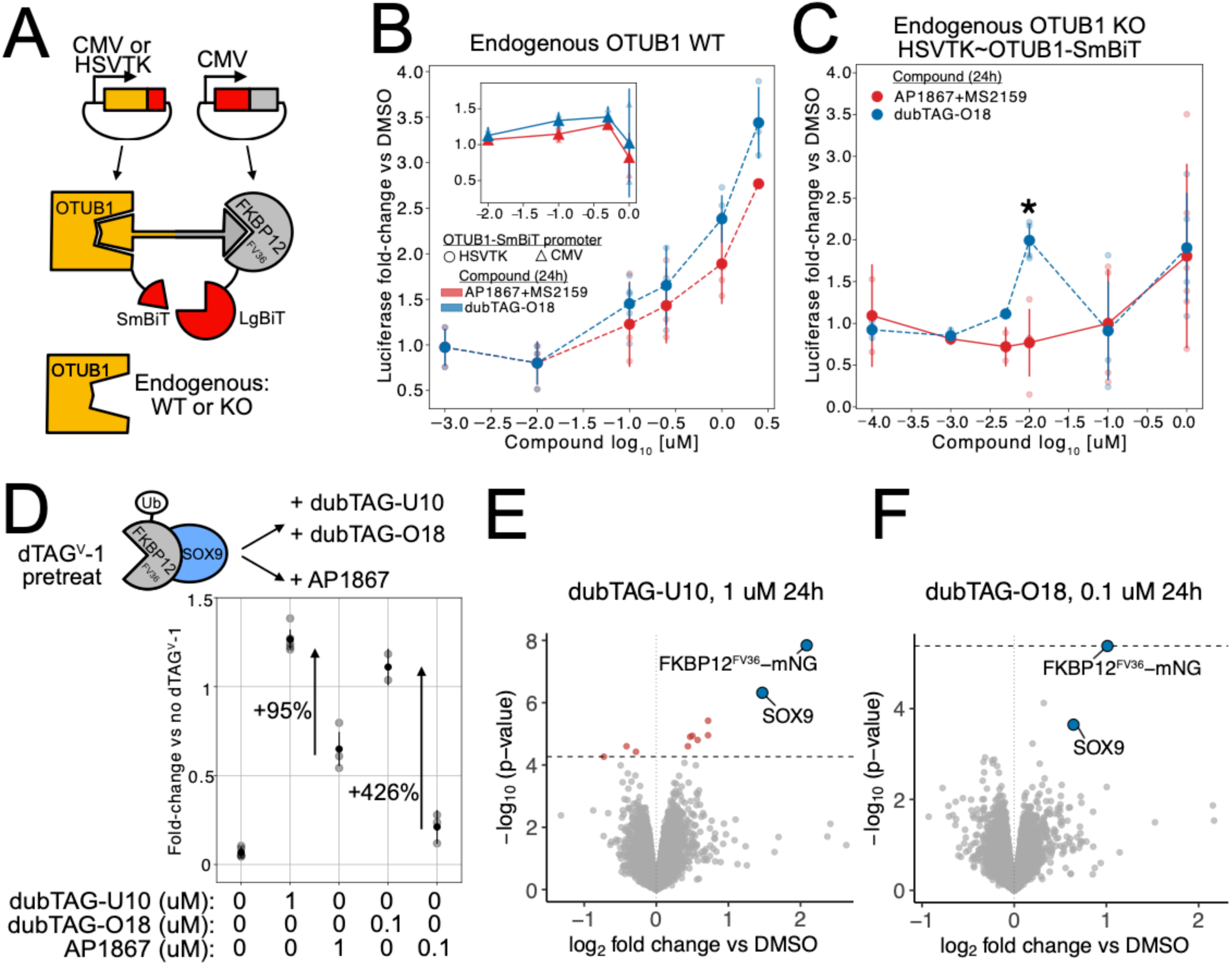
dubTAGs induce ternary complex formation and specifically increase FKBP12^F36V^-tagged protein levels through deubiquitinase recruitment. (A) Schematic of experiments using NanoBiT to demonstrate ternary complex formation in HEK293 cells. (B) Effect of dubTAG-O18 (blue) or binders (red) treatment on luciferase signal in cells expressing LgBiT-FKBP12^F36V^ and OTUB1-SmBiT under the control of weak HSVTK (outset, circles) or strong CMV (inset, triangles) promoter. (C) As in (B) but in HEK293 cells lacking endogenous OTUB1. *, p-value (two-sided t-test) < 0.01 comparing dubTAG-O18 to binders. One outlier with z-score > 2 was excluded. (D) Top, schematic of approach to assess deubiquitinase contribution to dubTAG activity by pretreatment with dTAG^V^-1 to destabilize SOX9 followed by comparison to FKBP12^F36V^ binder AP1867. Bottom, effect of indicated dubTAG or AP1867 treatments (all 24h) on SOX9 levels following 24h 5nM dTAG^V^-1 pretreatment. Light points represent independent biological replicates, dark points and error bars indicate mean and standard error. (E,F) Global proteomic effects (log_2_ fold change relative to DMSO) of dubTAG-U10 (E) or dubTAG-O18 (F) treatment after 24h 5 nM dTAG^V^-1 pretreatment. The dashed horizontal line indicates p-value corresponding to 5% FDR (n=3 biological replicates per condition).

We reasoned that these subtle effects could be due to high CMV-driven OTUB1 levels causing association with FKBP12^F36V^ even in the absence of dubTAG (i.e. at baseline). To test this, we expressed OTUB1-SmBiT under the control of the HSVTK promoter along with CMV-driven LgBiT-FKBP12^F36V^, resulting in a 54-fold reduction in baseline luciferase signal (Figure S3A), and repeated the NanoBiT assay. We observed a dose-dependent increase in luciferase signal with dubTAG-O18 treatment that was consistently higher than equimolar binders, culminating in a ∼3.5-fold increase at 2.5 μM (Figure 3B outset). However, this concentration was substantially higher than the 0.1 μM dubTAG-O18 that stabilized SOX9 in CNCCs and also induced a 2.7-fold change with binders. We hypothesized that these observations may reflect an excess of endogenous OTUB1 over the HSVTK-expressed OTUB1-SmBiT fusion. dubTAG-O18 would recruit relatively more of the endogenous OTUB1 to LgBiT-FKBP12^F36V^, so that only higher concentrations engage the less abundant OTUB1-SmBiT; at those same high concentrations, equimolar binders saturate endogenous OTUB1 and free OTUB1-SmBiT to form ternary complexes. This model predicts that performing NanoBiT in a context lacking endogenous OTUB1 should result in a) lower required dubTAG concentration and b) reduced binder effect. To test this prediction, we knocked out endogenous OTUB1 by CRISPR/Cas9 (Figure S3B) and repeated the NanoBiT assay. We observed the maximal effect (2-fold change in luciferase signal) with 10 nM dubTAG-O18, as compared to 2.5 μM in WT cells, and no discernable binder effect at 10 nM (Figure 3C). Together, these data indicate that dubTAGs induce ternary complex formation between their cognate deubiquitinases and FKBP12^F36V^ while also highlighting how concentrations of both exogenously expressed and endogenous targets can have substantial effects in the commonly used NanoBiT assay.

### dubTAG effects on target protein stabilization are specific and deubiquitinase recruitment-dependent

We next sought to test if the observed effects of dubTAG-U10 and dubTAG-O18 on SOX9 levels are driven by deubiquitinase-recruiting activity by comparing them to the FKBP12^F36V^ binder AP1867. We reasoned that conducting these experiments in a context where SOX9 is heavily destabilized through ubiquitination would result in a larger effect of deubiquitination and allow for a robust comparison between dubTAGs and AP1867. We therefore pre-treated SOX9-tagged CNCCs with the VHL-recruiter dTAG^V^-1 for 24h at 5 nM, reducing SOX9 levels to 5-8% of its normal levels (Figure 3D). We then continued dTAG^V^-1 treatment for 24h while adding dubTAG-U10 or dubTAG-O18 at their previously observed maximally effective concentrations (1 μM and 0.1 μM, respectively), comparing the resulting effects on SOX9 levels to equimolar AP1867 controls. Addition of dubTAG-U10 or dubTAG-O18 resulted in large increases in SOX9 levels (18 to 20-fold) from the low dTAG^V^-1 pre-treated baseline (Figure 3D), consistent with the idea that the effect of deubiquitinase recruitment on steady-state protein levels scales with the degree of pre-existing ubiquitination. Importantly, while equimolar AP1867 did result in some increase in SOX9 levels from the dTAG^V^-1 pre-treated baseline, presumably due to simple competition with dTAG^V^-1, both dubTAG-U10 and dubTAG-O18 showed significantly larger increases (95% and 426%, respectively) (Figure 3D). Treatment with a range of deubiquitinase binder (GNE-6776 or MS2159) concentrations showed minimal increases from the dTAG^V^-1 pre-treated baseline (Figure S3C). These results indicate that dubTAG-mediated effects on protein stability are potent and mediated by deubiquitinase recruitment.

To assess the specificity of dubTAG-U10 and dubTAG-O18, we performed 5 nM dTAG^V^-1 pre-treatment for 24h followed by addition of dubTAGs for 24h and assayed protein abundance changes by multiplexed, quantitative mass spectrometry-based proteomics . Across all samples, 9,780 proteins qualified for quantification, including peptides specific to SOX9 and the FKBP12^F36V^-mNeonGreen tag (Table S3). For dubTAG-U10 at 1 μM, differential protein expression analysis found 11 differential proteins at 5% FDR, of which SOX9 and FKBP12^F36V^-mNeonGreen showed the highest significance and fold-change magnitude (Figure 3E). Two of the other nine differentially expressed proteins (SYNPO and SPHK1) were found as SOX9-dependent genes in our previous study^12^, suggesting that they represent downstream effects of SOX9 upregulation rather than off-target effects. For dubTAG-O18 at 0.1 μM, FKBP12^F36V^-mNeonGreen was the only differentially expressed protein at 5% FDR, with SOX9 falling just below the threshold (Figure 3F). Consistent with our flow cytometry quantifications, dubTAG-U10 at 1 μM showed a greater increase in both SOX9 and FKBP12^F36V^-mNeonGreen abundance compared to dubTAG-O18 at 0.1 μM. These data confirm that dubTAGs selectively increase intracellular concentrations of FKBP12^F36V^ tagged proteins.

### Tunable and rapid reversal of dTAG-mediated degradation by dubTAGs

Having established that dubTAGs specifically stabilize FKBP12^FV36^-tagged proteins through deubiquitinase recruitment, we next asked if they could tunably reverse dTAG-mediated degradation to achieve distinct protein levels. Previous studies have relied upon washout of dTAG from cell culture media or cessation of *in vivo* delivery. While this approach has in some cases allowed for reversal of dTAG-mediated degradation^26,27^, it is not tunable and in some cases does not achieve full reversal^28^, likely due to its dependence on cellular division and dTAG off-rate. Indeed, applying this approach to SOX9-tagged CNCCs was unable to fully reverse dTAG^V^-1 mediated degradation: SOX9 levels peaked at approximately 38% of unperturbed levels 12h after washout of 5nM dTAG^V^-1 before settling at 20% by the 24h post-washout (Figure S4).

To assess whether dubTAGs can precisely reverse dTAG^V^-1-mediated degradation, we performed pre-treatment of SOX9-tagged CNCCs with 5 nM dTAG^V^-1 for 24h and then added dubTAG-U10, dubTAG-O18, or AP1867 at a range of concentrations for 24h. Across both dubTAG titrations, we observed up to nine distinct, reproducible SOX9 levels, ranging from 8% to over 120% of unperturbed SOX9 levels (Figure 4A). AP1867 alone only restored SOX9 to ∼60% of unperturbed levels even at 1 μM (Figure 4A), demonstrating the expanded range of restored protein levels enabled by deubiquitinase recruitment. Fitting Hill functions to the dose-response curves for dubTAG-U10 and dubTAG-O18 yielded similar EC_50_s (46.5 ± 2.9nM for U10 and 44.4 ± 4.3 nM for O18), indicating comparable effectiveness of the two molecules despite the slightly stronger effect size of dubTAG-U10 at higher concentrations.

**Figure 4.**
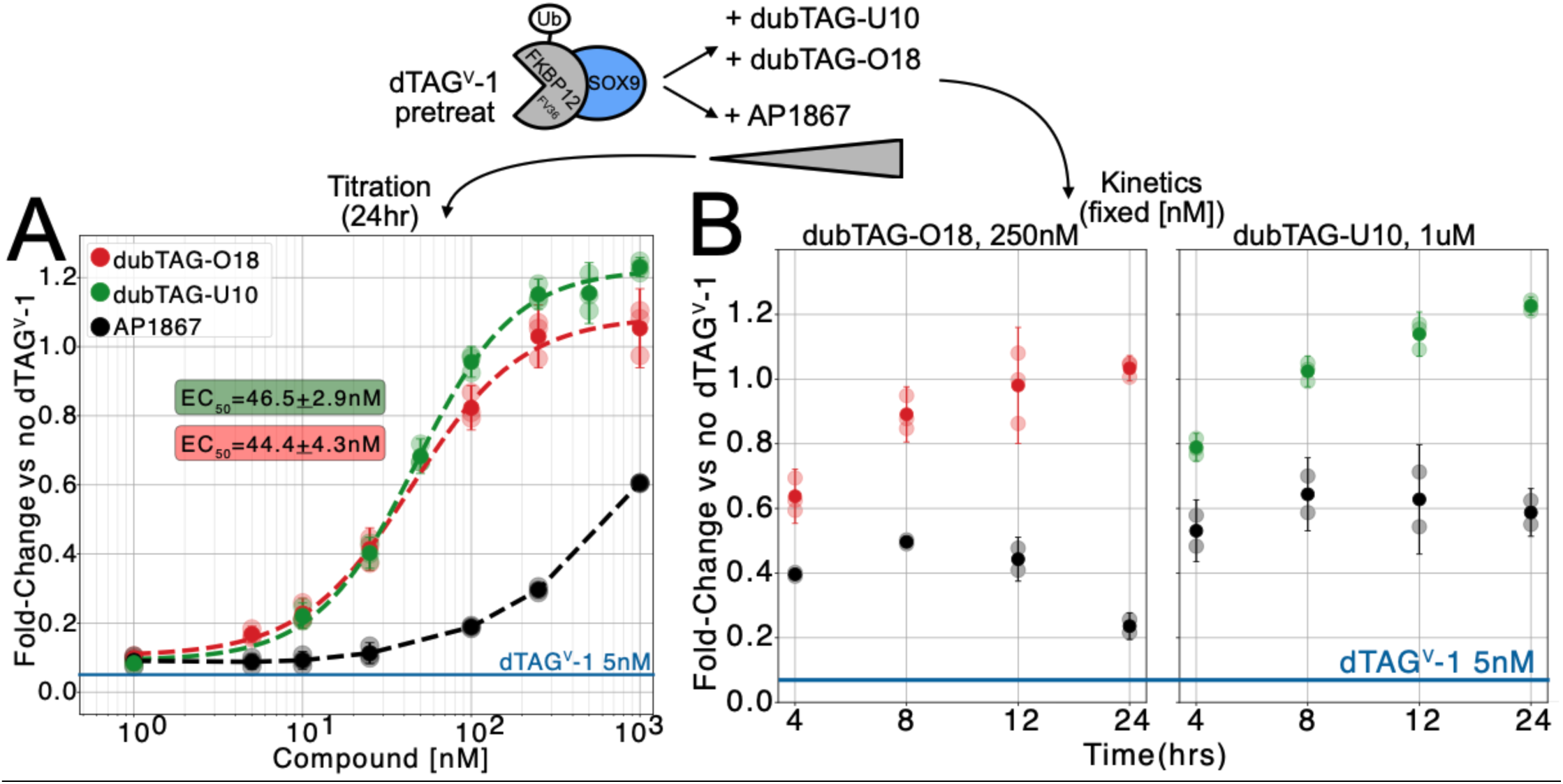
Tunable and rapid reversal of dTAG^V^-1-mediated degradation by dubTAGs. Top, approach to assess dose-dependence and kinetics of degradation reversal. (A) SOX9 levels as a function of dubTAG-U10 (green), dubTAG-O18 (red), or AP1867 (black) dose following 24h 5 nM dTAG^V^-1 pretreatment (mean indicated by blue line). Red and green dashed lines are Hill function fits, with EC_50_ values indicated +/-s.d. (B) SOX9 levels at indicated timepoints following application of dubTAG-O18 (left), dubTAG-U10 (right), or equimolar AP1867 (black) following 24h 5 nM dTAG^V^-1 pretreatment (mean indicated by blue line). Light points are independent biological replicates, dark points and error bars are mean and standard error.

We next assessed the kinetics of dubTAG reversal of protein degradation. We performed 24h of 5 nM dTAG^V^-1 pretreatment followed by dubTAG treatment fixed at the concentration that previously showed maximum dTAG^V^-1 reversal (1 μM for dubTAG-U10 and 0.25 μM for dubTAG-O18) and assayed SOX9 levels at 4h, 8h, 12h, and 24h timepoints. Both dubTAGs restored SOX9 levels to greater than 50% of unperturbed levels as early as 4h, and dubTAG-U10 completely restored SOX9 levels (100%) by 8h (Figure 4B). dubTAG-O18 restored SOX9 levels to within 10% of unperturbed levels by 8h at one-fourth the concentration of dubTAG-U10 and completely restored SOX9 levels by 12h. These effects were also driven by deubiquitinase recruitment, as equimolar AP1867 did not restore more than 70% of unperturbed SOX9 levels. Thus, our data demonstrate that dubTAGs enable tunable and rapid reversal of dTAG-mediated proteasomal degradation.

### Increasing SOX9 levels in CNCCs reveals both monotonic and non-monotonic responses in chromatin accessibility

We previously used dTAG^V^-1 to assess the effect of precise reductions in SOX9 levels on chromatin accessibility across the CNCC genome, finding a range of regulatory element (RE) responses ranging from highly sensitive to small reductions in SOX9 dosage to relatively buffered^12^. We therefore sought to use dubTAGs to understand the opposite end of the dosage spectrum, i.e. how increasing SOX9 above its wildtype levels in CNCCs impacts chromatin accessibility. We treated SOX9-tagged CNCCs with dubTAG-U10 for 48h, achieving two distinct dosages ∼25% and ∼50% above wildtype, and assessed genome-wide chromatin accessibility using ATAC-seq. We used dubTAG-U10 because it previously resulted in a larger increase in SOX9 levels compared to dubTAG-O18, and we chose a 48h timepoint to align with our previous work on reduced SOX9 dosage. We also included a full SOX9 depletion condition (500 nM dTAG^V^-1) to directly compare our newly generated data with data from our previous work. We first compared accessibility between all dubTAG-U10 and DMSO samples (four biological replicates each) at 151,457 reproducible ATAC-seq peak regions in CNCCs (which are candidate REs and are herein referred to as REs), finding 3,665 differentially accessible REs at 5% FDR. Of these, a roughly comparable number of REs showed gains (1,585) or losses (2,080) of accessibility (Figure 5A) (Table S4). Treatment with equimolar concentrations of AP1867 and the USP7 binder GNE-6776 found no differentially accessible REs at 5% FDR relative to DMSO (Figure S5A), indicating that the observed differential accessibility upon dubTAG treatment is driven by deubiquitinase recruitment and subsequent stabilization of SOX9. We next compared the changes in accessibility upon dubTAG treatment to changes resulting from full SOX9 depletion, integrating our previous SOX9 full depletion ATAC-seq to define a set of 46,559 REs differentially accessible upon either dubTAG-U10 treatment or full SOX9 depletion. We found a modest, positive correlation between the effect sizes for dubTAG-U10 treatment and full SOX9 depletion (adjusted R^2^ 0.249, Figure S5B), suggesting the existence of non-monotonic dosage responses to increased SOX9 levels.

**Figure 5.**
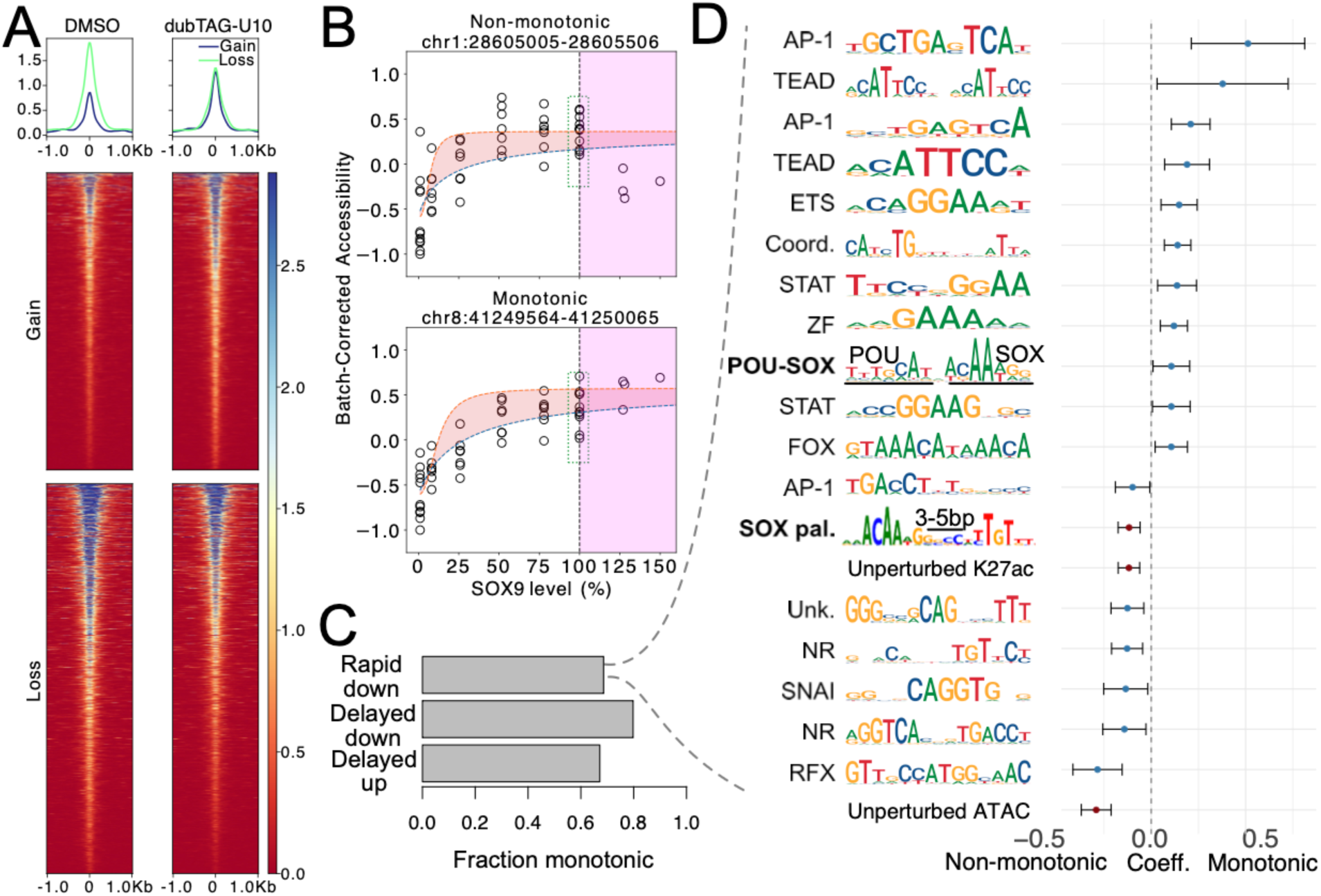
Increasing SOX9 levels in CNCCs reveals both monotonic and non-monotonic in chromatin accessibility driven by distinct regulatory features. (A) ATAC-seq signal from CNCCs treated with DMSO (left) or dubTAG-U10 (right), plotted over sets of REs with gained (n=1,585) or reduced (n=2,080) accessibility (5% FDR) following 48h treatment. (B) Schematic of method to distinguish non-monotonic (top) from monotonic (bottom) SOX9 dosage responses, with corresponding examples. Data in the pink area is from this study, data in white area is from Naqvi et al 2023 (plus additional replicates at 0% SOX9 level from this study). Shaded red curve represents 95% confidence interval fitting a Hill function to y-axis values at 100% and lower SOX9 dosage. (C) Fraction of REs showing monotonic responses as a function of time-dependence of SOX9 response. (D) Regression coefficients (x-axis, point is mean, error bars are 95% confidence intervals) of top 20 predictors of monotonic or non-monotonic among the rapidly responding (likely direct) SOX9 target REs. Consensus PWMs of motifs are displayed, and SOX motifs are bolded.

We next sought to systematically identify SOX9-dependent REs as monotonic or non-monotonic in the response to increased SOX9 levels. To do so, we integrated our newly generated ATAC-seq of dubTAG-U10 treatment with our previous ATAC-seq data titrating SOX9 levels downward with dTAG^V^-1. We found a strong correlation of full depletion effect with our previous study (adjusted R^2^ 0.81, Figure S5C). The presence of identical biological conditions (full SOX9 depletion and unperturbed samples) in both our newly generated dataset and the previous dataset allowed us to regress out technical confounders due to experimental batch and directly compare ATAC-seq counts per million (CPM) values along an expanded range of SOX9 dosage from ∼2% to 150% for each of 46,559 SOX9-dependent REs. We classified each of these REs as monotonic or non-monotonic by fitting a Hill function to the CPM values at SOX9 levels of 100% and lower, computationally extending this fitted curve above 100%, and finally testing whether the mean of the observed CPM values from SOX9 levels above 100% fell within the 95% confidence intervals of the partial Hill fit (see Methods for details) (Figure 5B) (Table S5). By this approach, 35,050 (75%) of the SOX9-dependent REs were monotonic and 11,449 (25%) were non-monotonic. We previously showed that SOX9-dependent REs are a mix of directly regulated targets with high SOX9 binding that respond rapidly (within 3h) of full SOX9 depletion and indirectly regulated targets that showed delayed responses^12^. The distribution of monotonic and non-monotonic responses was broadly similar between these distinct RE classes, with monotonic responses a majority (Figure 5C).

### SOX9 binding mode and interactions with other TFs drive monotonic and non-monotonic responses

Motivated by previous work showing that distinct sequence features drive buffered or sensitive responses to reduced SOX9 levels^24^, we sought to identify regulatory features associated with monotonic or non-monotonic SOX9 dosage responses. We focused on the subset of REs responding rapidly (within 3h) of full SOX9 depletion as these likely represent direct targets^12^. We assembled set of predictors for each of the 9,836 likely directly regulated REs; these predictors included the number of motif matches for each of 1,647 non-redundant TF motifs, as well as measured levels of both unperturbed chromatin accessibility and the active histone mark H3K27ac in CNCCs. We used regularized (least absolute shrinkage and selection operator [LASSO]) logistic regression to assess the contributions of these regulatory features to the probability of an individual RE showing a monotonic or non-monotonic response, yielding 23 predictors with non-zero coefficients (Table S6). The top predictor for non-monotonic responses was unperturbed chromatin accessibility level, with smaller contributions from RFX, and Class II nuclear receptor motif families as well as unperturbed H3K27ac levels.

Furthermore, the number of matches for a SOX palindrome sequence with 3-5 bp spacing, which we and others have demonstrated to be strongly and cooperatively bound by SOX9, was independently predictive of non-monotonic responses (Figure 5D). A diverse set of TF motifs emerged as predictive of monotonic RE responses, including motifs for the AP-1, TEAD, and ETS families as well as the previously described Coordinator motif ^23^. A composite POU-SOX motif consisting of a single POU site (TTTGCAT) and a single SOX site (ACAATG) with 1bp spacing was independently predictive for monotonic response (Figure 5D). This motif was most similar to the “canonical” POU-SOX motif that is cooperatively bound by POU5F1 (OCT4) and SOX2 in human and mouse stem cells. While previous *in vitro* DNA binding studies found minimal cooperativity between POU5F1 and SOX9 in binding to the composite POU-SOX motif^29^, POU5F1 is not expressed in CNCCs and POU3F3 and POU2F1 are instead the main expressed POU factors (Figure S6A). POU3F3 and POU2F1 diverge substantially from POU5F1 in amino acid sequence, including within the DNA binding domain, suggesting that these factors may be able to bind the composite motif through weak cooperativity with SOX9. Consistent with this idea, genomic instances of the POU-SOX motif showed significantly higher per-base contribution to SOX9 binding than random regions, as assessed by a BPNet sequence-to-activity deep learning model^30^ trained on SOX9 (V5) ChIP-seq. Both single SOX sites as well as the AP-1 motif showed no significant increase in contribution from random (Figure S6B). POU-SOX motif instances located within CNCC-accessible regions showed enrichment for SOX9 (V5) ChIP signal but significantly less signal than SOX palindrome motif instances; no such differences were observed in ChIP-seq from CNCCs lacking the V5 tag (Figure S6C). These results suggest that SOX9 binding mode and affinity combine with cross-TF interactions to drive divergent responses to increased SOX9 levels. More broadly, our results show how dubTAGs can be used to understand the regulatory effects of quantitatively increased TF levels, questions largely inaccessible by existing chemical genetic approaches.

## Discussion

We have developed and applied heterobifunctional small molecules (dubTAGs) that recruit cellular deubiquitinases to the FKBP12^F36V^ degron tag and stabilize endogenously tagged proteins in mammalian cells. Our goal was to complement dTAG and other targeted protein degradation systems by providing a means to rapidly and precisely increase proteins above their typical levels and reverse targeted protein degradation. Given the rapid adoption of dTAG to assess the immediate consequences of protein depletion, we anticipate that dubTAGs will provide a readily applicable tool to investigate consequences of either increased protein levels or rapid restoration of degraded protein on cellular processes.

An inherent limitation of our approach is that dubTAGs increase protein levels by reversing ubiquitination and resultant degradation, and therefore rely on baseline transcription and translation rates to accumulate protein. This should suffice for most proteins at steady-state and especially for reversing targeted degradation (where ubiquitination is by definition rate-limiting), but may not extend to proteins with low endogenous transcription or translation and/or long protein half-lives. Given that over 80 proteins have been endogenously tagged with FBKP12^F36V^ (Table S1) and that USP7 and OTUB1 (the endogenous deubiquitinases recruited by dubTAGs) are broadly expressed, broader application of dubTAGs should allow for the assessment of these distinct contributions.

Since the development of dTAG, others have developed degron systems that use orthogonal tags and binders, including bromoTag and HaloTag^31,32^. This opens up the exciting possibility of combinatorial, bidirectional protein dosage control, in which one protein of interest is tagged with FBKP12^F36V^ and modulated with dTAG or dubTAGs and the other is tagged with bromoTag or HaloTag and modulated with an orthogonal set of heterobifunctional molecules. Indeed, a recent study demonstrated stabilization of the lysosomal enzyme PRCP when it was exogenously expressed and tagged with HaloTag^31–33^; future studies should extend and validate such approaches for endogenous protein targets, as we have done here for dubTAGs.

We previously used dTAG^V^-1 to study the effect of reduced SOX9 levels on chromatin accessibility in cranial neural crest cells (CNCCs), finding responses of individual regulatory elements (REs) ranging from buffered to highly sensitive. Here, using dubTAGs to stabilize SOX9 above wild-type levels, we find a bifurcated chromatin accessibility response: most REs change monotonically with SOX9 dosage, but a sizable minority show non-monotonic responses. These divergent responses are driven by distinct sequence features: non-monotonic responses are associated with high-affinity SOX palindrome motifs, whereas monotonic responses are in part associated with a composite POU-SOX motif. A potential explanation for these results is that increasing SOX9 levels via dubTAG stabilization allows SOX9 dimers to form off of DNA; these unproductive dimers cannot bind the SOX palindrome in the canonical sequential manner, producing non-monotonic accessibility at those sites. Composite POU-SOX motifs, by contrast, may function as low-affinity monomeric SOX9 binding sites and can therefore continue to gain binding and yield monotonic accessibility responses with increasing SOX9 levels. Function of the composite POU-SOX motif has been previously demonstrated in stem cells when bound by SOX2/17 and POU5F1 (OCT4)^34,35^; our data suggest it may be engaged by SOX9 with other POU factors (POU3F3/POU2F1), highlighting how dubTAGs can reveal novel insights into TF biology.

Although our primary application of dubTAGs in this work was to assess the effect of increasing TF levels above 100%, we also demonstrate that dubTAGs can be used to precisely and rapidly reverse dTAG-mediated degradation (Figure 4). This functionality could be used to assess the reversibility of chromatin and gene expression effects induced by TF degradation and dissect the dosage-dependence and kinetics of this reversibility. More broadly, dubTAG-mediated restoration of protein levels could be used to investigate cellular “memory” for any protein of interest and its dependent cellular processes.

## Methods

### dubTAG synthesis

All commercial chemical reagents and solvents were used for the reactions without further purification. Flash column chromatography was performed on Teledyne ISCO CombiFlash Rf+ instrument equipped with a 220/254/280 nm wavelength UV detector and a fraction collector. Normal phase column chromatography was conducted on silica gel columns with either hexane/ethyl acetate or dichloromethane/methanol as eluent. Reverse phase column chromatography was conducted on HP C18 RediSep Rf columns, and the gradient was set to 10% of acetonitrile in H_2_O containing 0.1% TFA progressing to 100% of acetonitrile. All final compounds were purified with preparative high-performance liquid chromatography (HPLC) on an Agilent Prep 1290 infinity II series with the UV detector set to 220/254/280 nm at a flow rate of 40 mL/min. Samples were injected onto a Phenomenex Luna 750 x 30 mm, 5 μm C18 column, and the gradient was set to 10% of acetonitrile in H_2_O containing 0.1% TFA progressing to 100% of acetonitrile. LCMS was performed by an Agilent 1200 series system with DAD detector and a 2.1 mm x 150 mm Zorbax 300SB-C18 5 μm column for chromatography and high-resolution mass spectra (HRMS) that were acquired in positive ion mode using an Agilent G6230BA Accurate Mass TOF with an electrospray ionization (ESI) source. Samples (0.8 μL) were injected onto a C18 column at room temperature, and the flow rate was set to 0.6 mL/min with water containing 0.1% formic acid as solvent A and acetonitrile containing 0.1% formic acid as solvent B. Nuclear magnetic resonance (NMR) spectra were acquired on Bruker DRX 400 MHz for proton (^1^H NMR) and 101 MHz for carbon (^13^C NMR). Chemical shifts for all compounds are reported in parts per million (ppm, δ). The format of chemical shift was reported as follows: chemical shift, multiplicity (s = singlet, d = doublet, t = triplet, q = quartet, m = multiplet), coupling constant (J values in Hz), and integration. All final compounds had > 95% purity using the HPLC methods described above, and were in trifluoroacetate salt form. Please refer to the supporting information for detailed synthetic routes, procedures, and data.

### hESC culture and differentiation

Wildtype or SOX9-tagged hESC lines were cultured on wells coated with Matrigel (Corning-354277) and passaged with ReLeSR (StemCell Technologies) every 4-6 days depending on initial seeding and overall colony density. Differentiation of hESCs was performed according to a protocol derived from Bajpai et al, Prescott et al, and Long et al ^16,36,37^. Briefly, hESC colonies were dissociated with Type IV collagenase (StemCell Technologies-7426) and plated in a low-attachment dish, in differentiation media (1:1 ratio DMEM F-12 and Neurobasal supplemented with a final concentration 0.5x Glutamax, 0.5x N2 Supplement (StemCell Technologies - 7152), 0.5x B27 (StemCell Technologies - 5711), 20ng/mL of rFGF (Fisher - BT-FGFB-100), 20ng/mL of EGF (StemCell Technologies-78006.1) and 5ug/mL of Bovine Insulin (Fisher-NC2011227).

Between 9-11 days in differentiation media, embryoid bodies attach to the dish and immature neural crest cells begin to migrate from these attachments. The CNCCs were then passaged into Fibronectin (7.5ug/mL)-coated plates at a density of approximately 1.56x10^5 cells/cm^2 and treated with CNCC Maintenance media (same as the above-described differentiation media with the exception of swapping Bovine Insulin with 1mg/mL of BSA). At passage 2, CNCCs were transitioned to Maintenance media supplemented with 1ng/mL of BMP-2 (StemCell Technologies - 78004) and 3 μM of Wnt Agonist CHIR 99021 (StemCell Technologies - 72054). For all experiments, we used either passage 4 or 5 CNCCs, seeding them into 24, 48, or 96 well plates depending on cell density requirements.

### Flow cytometry measurements of FKBP12^F36V^ tagged TFs

To screen dubTAGs in hESCs, SOX9-tagged hESCswere allowed to grow for 4 to 6 days from passage and then dissociated to single cells using StemPro Accutase (Gibco-A1110501). Cells were plated at a density of 6.8x10^4 cells/cm^2 in mTESR Plus media supplemented with 10 μM Y-27632 (StemCell Technologies - 72304). dubTAGs were applied at a concentration of 1 μM over a 48hr treatment window, with treatment media changed every 24h. Wells were then washed with PBS, treated with Accutase for 10-15 minutes until single cell suspensions were formed and stained with the 7aaD viability dye (Biologened-420404). SOX9 levels were quantitatively measured using flow cytometry, gating cells based on viability (Per-CP channel) to measure mNeonGreen (FITC) expression. Wildtype (untagged, lacking mNeonGreen) hESCs were used as a negative control.

To assess the effects of the dubTAGs in SOX9-tagged CNCCs, passage 4 or 5 CNCCs were plated at densities ranging from 6.3 to 9.4 x10^4 cells/cm^2. After overnight incubation, cells were applied with treatment media for 48h. For assays evaluating kinetics and dosage of dTAG^v^-1 induced SOX9 degradation recovery, 5nM dTAG^v^-1 was applied to treatment wells for a 24h followed by a combination of 5nM dTAG^V^-1 treatment and varying concentrations of dubTAGs for an additional 24h. To screen the effects of multiple candidate dubTAGs across concentration and time, CNCCs were cultured in either 48-well or 96-well plates and treated at the required concentration and duration. Cells were then treated with Accutase for 8 minutes to ensure single cell suspensions before staining with 7-AAD viability dye and transferred to 96-well plates for high-throughput measurement of mNeonGreen (SOX9) using the BD LSRFortessa X-20 plate reader system with the FITC and Per-CP channels to assay SOX9 expression levels and gate cells for viability, respectively.

To measure dubTAG effects on TWIST1 levels, passage 4 TWIST1-tagged CNCCs were cultured in either 6-well or 12-well plates and treated for 48h with specific dubTAGs. CNCCs were dissociated from wells with Accutase and fixed with 4% PFS, followed by permeabilization with 0.1% Triton-X for an hour. Cells were then blocked in a 1% BSA solution with goat serum at 3% w/v for 45 minutes and stained with the primary antibody for the V5 epitope (ThermoFisher, #R960-25) (1:200 dilution) for an hour. Cells were washed twice with 1x PBS before secondary staining with Alexa 488 tagged goat-anti mouse IgG secondary (ThermoFisher, #A-11001) for an hour. After three additional PBS washes, we measured TWIST1 levels using flow cytometry. The .fcs acquisition files containing all channel information from flow cytometry runs were processed via custom scripts in Python to calculate the effect of different treatment regimes on FKBP^F36V^ tagged TF levels.

### NanoBiT assay

#### Cloning and Expression of OTUB1-SmBit and LgBiT-FKBP12^F36V^ fusion constructs

Plasmids expressing the split NanoBiT luciferase gene pcDNA3.1-LgBiT-G(SSS) Linker and pcDNA3.1-G(SSS) Linker-SmBiT (referred to as LgBiT and SmBiT plasmids) were a gift from Dr. Jonathan Kagan. We designed primers to amplify the coding region of OTUB1 from FLAG-OTUB1 WT-pcDNA3.1 (gift from Cynthia Wolberger, Addgene #118209) and FKBP12^F36V^ from pAAV-hSOX9-dTAG-mNeonGreen-V5 ^12^, with homologous sequences specific to insertion sites in the SmBiT and LgBiT plasmids respectively along with primers to amplify the vectors. PCR amplifications were performed using Q5 High-Fidelity 2X Master Mix (NEB M0492S). Final constructs were synthesized by assembling the backbone vector and insertion sequences using the NEBuilder HiFi DNA Assembly Master Mix (NEB E2621S) and transformed into chemically competent DH5 alpha cells (NEB C2987H). Single colonies were validated by restriction enzyme digests for insertion of OTUB1 and FKBP12^F36V^ into the NanoBiT plasmid vectors and verified by complete plasmid sequencing. To generate HSVTK-FLAG-OTUB1-G(SSS)Linker-SmBiT, we sub-cloned the HSVTK promoter into our assembled pcDNA3.1 OTUB1-G(SSS) Linker-SmBiT from psiCheck2-SV40p-3xFLAG-HSVTKp-mCherry (gift from Chunghun Lim, Addgene #182377).

### Transfection and Measurement of NanoBiT Plasmids

Transfection of OTUB1-SmBiT and LgBiT-FKBP-expressing plasmids into HEK293 cells was performed using Lipofectamine 2000 (Invitrogen 11668030) and was optimized for individual reactions in a 96-well plate format. Each reaction used a total of 200ng of DNA split in a 1:1 molar ratio between the OTUB1-smBiT and LgBiT-FKBP plasmids. Lipofectamine 2000 reagent was initially added to DMEM-HG (Cytivia SH30081) at a ratio of 1(ug DNA):3(ul Lipofectamine) and allowed to incubate for 5 minutes. The DNA and Lipofectamine tubes were then combined and allowed to sit at 20 minutes at room temperature before aliquoting 50ul of transfection complex per-well of a 96-well plate followed by addition of HEK 293 cells.

HEK293 cells were cultured in 293 complete media (1x NEAA, 1x Glutamax, 1x Sodium Pyruvate, 10% FBS) with a seeding density of 3.125x10^4 cells/cm^2, and passaged every 2-3 days depending on confluency. On the day of transfection, cells were treated with 0.25% Trypsin-EDTA for 5 minutes at 37 degrees to dissociate cells and quenched with 293 complete media post-incubation. Cells were then triturated with a P1000 to generate single cell suspensions, collected in a 15ml conical, centrifuged at 300xg for 4 minutes and resuspended in 293 complete media. After counting, cells were plated into a 96-well plate with a seeding density of 50,000 cells/well directly into the transfection complex. Media was changed approximately 24 hr after the start of incubation for treatment with different concentrations of dubTAG-O18 or the control binder (equimolar amounts of MS2159 and AP1867) for a further 24h. At the end of treatment, flurofurimazine (VWR A1355226) - the substrate for the NanoLuc Enzyme - was added to the wells at a final concentration of 4uM and luminescence was measured on a TECAN Spark.

### Generation of OTUB1 knockout HEK293 cells

OTUB1 knockout HEK293 cells were generated using CRISPR-Cas9 gene editing. Plasmid PX459, expressing both the Cas9 nuclease and sgRNA (Addgene #62988)^38^, was used to subclone 4 different sgRNAs targeting the OTUB1 gene (see ^39^ for OTUB1 sgRNA sequences, AAVS1 sgRNA was sequence was 5’-GTTAATGTGGCTCTGGTTCT-3’). sgRNAs were ordered as separate oligos with complementary and reverse complement sequences prior to annealing and subcloned into PX459 by sequential digestion and ligation with BbsI Type IIS restriction enzyme (NEB R3539S) at 37 degrees and T4-DNA ligase (NEB M0202S) at 16 degrees in 5 minute intervals for 30 cycles. Reactions were transformed into NEB Stable cells (NEB C3040H) and incubated at 30 degrees for 24h. Single colonies of sgRNA subcloned into PX459 were verified by full plasmid sequencing.

For transfection of OTUB1-or AAVS1-sgRNA plasmids, early passage HEK 293 cells were seeded in a 12-well plate at a density of 9.4x10^4 cells/cm^2 and were between 70-90% confluency on the day of transfection. Transfection complexes were formed using Lipofectamine 2000, with a 1:5 DNA to Lipofectamine ratio and transfected according to the Manufacturer’s protocol. After 24h, cells were passaged and seeded at a density of 0.9x10^6 cells/well in a 6-WP with 293 complete media supplemented with 1mg/ml of Puromycin for selection. Puromycin treatment was carried out for 72h, after which cells were allowed to recover and further passaged for harvesting cell lysate and cryopreservation. Total soluble protein lysate was quantified using BCA Protein Assay Kit (Thermo 23227), and equal amounts of lysate were separated on an SDS-page gel and subject to Western Blot with anti-OTUB1 (Cell Signaling Technologies 3783T) at 1:1000 dilution, with Goat anti-Rabbit HRP antibody (vendor) at 1:2000 dilution. HRP-conjugated Beta-actin (Biolegend 643807) at 1:5000 dilution was used as a loading control. HEK293 cells transfected with OTUB1 sgRNA 2 were chosen for NanoBit analysis.

### ATAC-seq collection and library transposition

SOX9-tagged CNCCs seeded in 24-Well Plate with CNCC Maintenance media supplemented with BMP-2 and CHIR99021. After overnight incubation, cells were treated with 1 μM of dubTAG-U10 or vehicle control for a total treatment of 48h - with fresh treatment media applied every 24h. As the dubTAG-U10 consists of the FKBP12^F36V^ binder AP1867 and the USP7 binder GNE-6776, we treated cells with a binder control consisting of 1 μM of each molecule to rule out the possibility that accessibility changes from dubTAG-U10 were driven by altered functionality of SOX9-FKBP12^F36V^ and changes to USP7 structure or function. Post treatment, media was removed, cells were washed with PBS once, and then treated with accutase (50ul per well) to generate single cell suspensions. Cells across treatment wells were harvested and counted, with 6x10^4 cells portioned for transposase reactions. Transposition was performed using Illumina Tagmentation transposase master mix (Active Motif - 53152) according to the method of Grandi et al ^40^. Post transposition, sample transposed DNA was amplified using Illumina i5 and i7 indexes to generate libraries and quantified using rt-QPCR to determine the appropriate number of additional PCR cycles to reach a target library concentration of 240 fmol. Sample libraries were then pooled and shipped to Novogene for Paired-End 150 bp sequencing.

### Sample preparation for mass spectrometry (total proteome profiling)

CNCCs were plated at a density of 650,000 cells per-well on fibronectin coated wells of a 6-Well plate (final Fibronectin concentration: 7.5ug/mL). After overnight incubation, treatment wells were pre-treated with 5nM dTAG^V^-1 for 24h to reduce SOX9-FKBP12^F36V^-mNeonGreen levels, after which treatment media was applied for an additional 24h. Treatment media consists of 5nM dTAG^V^-1 along with either 0.1 μM of dubTAG-018 or 1 μM of dubTAG-U10. Cells were immediately washed 3-times in Ice-cold PBS (2mL wash per well) after treatment, followed by final application of 450ul of ice-cold PBS across wells. Samples were then placed on ice, and cells were scraped off the plate and transferred to eppendorf tubes. Samples were promptly flash-frozen in liquid nitrogen and stored at -80 degrees for approximately a week before sample preparation. Samples for protein analysis were prepared essentially as previously described ^41^. Proteomes were extracted using a buffer containing 200 mM EPPS pH 8.5, 8M urea, 0.1% SDS and protease/phosphatase inhibitors. Following lysis, 50 µg of each proteome was reduced with 5 mM TCEP. Cysteine residues were alkylated using 10 mM iodoacetimide for 20 minutes at RT in the dark. Excess iodoacetimide was quenched with 10 mM DTT. A buffer exchange was carried out using a modified SP3 protocol^42^. Briefly, ∼500 µg of Cytiva SpeedBead Magnetic Carboxylate Modified Particles (65152105050250 and 4515210505250), mixed at a 1:1 ratio, were added to each sample. 100% ethanol was added to each sample to achieve a final ethanol concentration of at least 50%. Samples were incubated with gentle shaking for 15 minutes. Samples were washed three times with 80% ethanol. Protein was eluted from SP3 beads using 200 mM EPPS pH 8.5 containing Lys-C (Wako, 129-02541). Samples were digested overnight at room temperature with vigorous shaking. The next morning trypsin (ThermoFisher Scientific) was added to each sample and further incubated for 6 hours at 37° C. For experiments that were split between plexes, a bridge sample was generated by pooling a small volume of each digest. This pooled sample was then included as a 17th sample in each of the plexes. Acetonitrile was added to each sample to achieve a final concentration of ∼33%. Each sample was labelled, in the presence of SP3 beads, with ∼125 µg of TMTPro reagents (ThermoFisher Scientific). Following confirmation of satisfactory labelling (>97%), excess TMT was quenched by addition of hydroxylamine to a final concentration of 0.3%. The full volume from each sample was pooled and acetonitrile was removed by vacuum centrifugation for 1 hour. The pooled sample was acidified and peptides were de-salted using a Sep-Pak 50mg tC18 cartridge (Waters). Peptides were eluted in 70% acetonitrile, 1% formic acid and dried by vacuum centrifugation.

### Basic pH reversed-phase separation (BPRP)

TMT labeled peptides (phosphopeptide enrichment flow through from phosphoproteomics experiment) were solubilized in 5% acetonitrile/10 mM ammonium bicarbonate, pH 8.0 and ∼300 µg of TMT labeled peptides were separated by an Agilent 300 Extend C18 column (3.5 m particles, 4.6 mm ID and 250 mm in length). An Agilent 1260 binary pump coupled with a photodiode array (PDA) detector (Thermo Scientific) was used to separate the peptides. A 45 minute linear gradient from 10% to 40% acetonitrile in 10 mM ammonium bicarbonate pH 8.0 (flow rate of 0.6 mL/min) separated the peptide mixtures into a total of 96 fractions (36 seconds). A total of 96 Fractions were consolidated into 24 samples in a checkerboard fashion and vacuum dried to completion. 12 of the 24 samples were desalted via Stage Tips and re-dissolved in 5% formic acid/ 5% acetonitrile for LC-MS3 analysis.

### Liquid chromatography separation and tandem mass spectrometry (LC-MS3)

Proteome data were collected on an Orbitrap Eclipse mass spectrometer (ThermoFisher Scientific) coupled to a Proxeon EASY-nLC 1000 LC pump (ThermoFisher Scientific). Fractionated peptides were separated using a 120 min gradient at 550 nL/min on a 35 cm column (i.d. 100 μm, Accucore, 2.6 μm, 150 Å) packed in-house. A FAIMS device enabled during data acquisition with compensation voltages set as −40, −60, and −80^43^. MS1 data were collected in the Orbitrap (60,000 resolution; maximum injection time 50 ms; AGC 4 × 105). Charge states between 2 and 5 were required for MS2 analysis in the ion trap, and a 120 second dynamic exclusion window was used. Top 10 MS2 scans were performed in the ion trap with CID fragmentation (isolation window 0.5 Da; Turbo; NCE 35%; maximum injection time 35 ms; AGC 1 × 104). Real-time search was used to trigger MS3 scans for quantification^42,44^. MS3 scans were collected in the Orbitrap using a resolution of 50,000, NCE of 55%, maximum injection time of 250 ms, and AGC of 1.25 × 105. The close out was set at two peptides per protein per fraction.

### Mass spectrometry data analysis

Raw files were converted to mzXML, and monoisotopic peaks were re-assigned using Monocle^45^. Searches were performed using the Comet search algorithm against a human database downloaded from Uniprot in May 2021. We used a 50 ppm precursor ion tolerance, 1.0005 fragment ion tolerance, and 0.4 fragment bin offset for MS2 scans collected in the ion trap, and 0.02 fragment ion tolerance; 0.00 fragment bin offset for MS2 scans collected in the Orbitrap. TMTpro on lysine residues and peptide N-termini (+229.1629 Da) and carbamidomethylation of cysteine residues (+57.0215 Da) were set as static modifications, while oxidation of methionine residues (+15.9949 Da) was set as a variable modification. Each run was filtered separately to 1% False Discovery Rate (FDR) on the peptide-spectrum match (PSM) level. Then proteins were filtered to the target 1% FDR level across the entire combined data set ^45,46^. For reporter ion quantification, a 0.003 Da window around the theoretical m/z of each reporter ion was scanned, and the most intense m/z was used. Reporter ion intensities were adjusted to correct for isotopic impurities of the different TMTpro reagents according to manufacturer specifications. Peptides were filtered to include only those with a summed signal-to-noise (SN) ≥ 90 across all TMT channels. The signal-to-noise (S/N) measurements of peptides assigned to each protein were summed (for a given protein). These values were normalized so that the sum of the signal for all proteins in each channel was equivalent thereby accounting for equal protein loading. Differential protein expression testing was performed with the DEqMS package (v3.23) ^47^, accounting for CNCC differentiation batch as a covariate. BH method-adjusted DEqMS p-values were used for FDR control.

### ATAC-seq data processing

Nextera adapter sequences and low-quality bases (-Q 10) were trimmed from sequencing reads using skewer v0.2.2 ^48^ and aligned to the human genome (hg38) using bowtie2 v2.4.1^49^ with the following settings: --very-sensitive, --X 2000. Read mate pair information was corrected with samtools v1.10 fixmate, PCR duplicates were removed using samtools markdup, and mitochondrial reads and low-mapping-quality reads (-q 20) were removed using samtools v1.10 view^50^. Shifted bed sites representing Tn5 insertions were obtained from mapped and filtered ATAC bam files and for each library, insertions were counted over a set of 151,457 reproducible peak regions from^12^, which are candidate regulatory elements and referred to as REs.

### Differential accessibility testing and visualization

To determine differentially accessible REs in CNCCs treated with dubTAG-U10 at 1 μM, dTAG^V^-1 at 500nM, or the control binder (AP1867 and GNE-6776 both at 1 μM) relative to vehicle treatment, we analyzed Tn5 insertion counts over the set of 151,457 REs generated from the ATAC-seq data processing pipeline mentioned above. We augmented the datasets with the vehicle treatment and full SOX9 depletion (500nM dTAG^V^-1) samples from our previous work^12^ and filtered the combined dataset to include only REs with at least 100 counts across all samples. We then used the DeSeq command from the R package DEseq2^51^ to assess the mean and variance of the modeled binomial distribution of gene counts per sample using treatment condition, differentiation batch, and edited cell line (two clonal lines of SOX9-tagged CNCCs were used in this study) as covariates. We then called differentially accessible REs at a 5% FDR between conditions (Vehicle vs 1 μM dubTAG-U10 or Vehicle vs 500 nM dTAG^V^-1). For display of differentially accessible REs from dubTAG U-10 treatment as genomic heat maps, we used the macs2 peak calling algorithm^52^ to generate narrow peaks from the resulting shifted bed files (Tn5-corrected peak coordinates) of the ATAC-seq data processing, selecting one representative sample from the dUBTAG-U10 and DMSO control condition. We stratified genomic regions based on overlap with REs that showed differential loss or gain of accessibility and retrieved these coordinates from the narrow peak .bed files using the bedtools intersect command. We then used the deepTools package^53^ to generate the matrix file for counts across the regions between the representative samples with the computeMatrix reference-point command using a 1kb window around the peak center and a bin-size of 25bp. To plot the heat-map of the differentially accessible REs, we used the deepTools plotHeatMap command.

### Modeling of monotonic and non-monotonic SOX9 dose-response curves

To evaluate whether REs exhibited non-monotonic profiles at higher SOX9 levels resulting from dubTAG-U10 treatment, we integrated our previous ATAC-seq data titrating SOX9 levels downward with dTAGV-1 (five concentrations) with the chromatin accessibility data from 1 μM dubTAG-U10-treated SOX9-tagged CNCCs, yielding a total of eight distinct SOX9 levels. We normalized the integrated dataset using the trimmed mean of log-expression ratios^54^ to generate counts per million values for each RE and used the lm function in R to regress out differentiation batch and clonal line, removing batch effects and non-treatment contributions. We then fit a logistic equation to the batch-corrected RE counts per million for the set of 46,559 differentially accessible REs using the drc package in R across the SOX9 concentration range at or below endogenous levels. For each RE, we determined whether a 2-parameter logistic equation (free parameters: signed Hill coefficient and ED50) or a 3-parameter fit (free parameters: signed Hill coefficient, ED50, and max signal) optimally fit the data, accepting the 3-parameter fit if the difference in AIC values between the models was at least 2.

Applying confint(), we determined the upper and lower 95% confidence intervals for each fitted parameter to compute boundary logistic functions across the dubTAG-U10 treatment range (100–150% SOX9). Finally, we evaluated the mean batch-corrected counts per million for a given RE at SOX9 levels >100% against the fitted boundary conditions: if the mean value fell below the lower bound for an RE with a negative signed Hill coefficient (batch-corrected accessibility increases from depleted SOX9 levels to 100%), we assigned that RE as non-monotonic. For REs with a positive signed Hill coefficient (accessibility increases as SOX9 is depleted), we assigned non-monotonic status if the mean exceeded the upper bound of the fitted logistic function.

### Sequence analysis of monotonic and non-monotonic REs

Full RE sequences were scanned for motif matches using the BagOfMotifs R package (runFIMO command with default parameters)^55^ using a set of non-redundant motifs from the GimmeMotifs vertebrate v5.0 motif database^56^. For each RE, the number of matches for each motif (permissive q-value < 0.5) was counted, and additional predictors including baseline ATAC-seq counts, baseline H3K27ac counts, CpG counts, GC content, and number of SOX palindrome motif matches were added (using data from^12^). To select informative motif features, we fit an L1-regularized (LASSO) logistic regression using glmnet with 10-fold cross-validation, using the above-described set of predictors to predict monotonic versus non-monotonic class. Features with non-zero coefficients at lambda.1se were retained. We then refit a standard (unregularized) logistic regression on the selected features to obtain interpretable coefficient estimates, standard errors, and 95% confidence intervals (coefficient ± 1.96 × SE). Features were ranked by absolute coefficient magnitude.

## Data availability

The mass spectrometry proteomics data have been deposited to the ProteomeXchange Consortium via the PRIDE partner repository with the dataset identifier PXD082352; processed peptide counts are available on Zenodo (https://zenodo.org/records/22132638). Raw ATAC-seq data generated during this study will be made available on the Gene Expression Omnibus (accession number GSE345440); processed ATAC-seq counts for REs are available on Zenodo (https://zenodo.org/records/22129390). Plasmids generated in this study will be deposited in Addgene. All other reagents are available upon request to S.N. or J.J.

## Code availability

Custom code for analyzing processed ATAC-seq and proteomics data is available on Zenodo (https://zenodo.org/records/22132638).

## Acknowledgements

We acknowledge the Boston Children’s Hospital Flow Cytometry and the Thermo Fisher Scientific Center for Multiplexed Proteomics at Harvard Medical School (https://tcmp.hms.harvard.edu) for quantitative proteomics. This work was supported in part by the endowed professorship from the Icahn School of Medicine at Mount Sinai (to J.J.) and utilized the NMR Spectrometer Systems at Mount Sinai acquired with funding from the National Institutes of Health (NIH) SIG Grants 1S10OD025132 and 1S10OD028504. S.N. acknowledges support from the Charles H. Hood Foundation, the Richard and Susan Smith Family Foundation, and NIH Grants R35GM165577 and R00DE032729.

## Author contributions

Conceptualization, S.G, W.W, J.J, S.N.; methodology, S.G., S.S., and X.S.; software, S.G. and S.N.; formal analysis, S.G. and S.N.; investigation, S.G. and S.S.; resources, S.G., X.S., Q.W., J.J., Y.X., and S.N.; writing – original draft, S.G. and S.N.; writing – review & editing, S.G., S.N., Y.X., X.S.,, and J.J.; visualization, S.G. and S.N.; supervision, S.N., Y.X., and J.J.; project administration, S.N., Y.X., and J.J.; funding acquisition, S.N., and J.J.

## Notes

The authors declare the following competing financial interest(s): The Jin laboratory received research funds from Celgene Corporation, Levo Therapeutics, Inc., Cullgen, Inc. and Cullinan Therapeutics, Inc. J.J. is an equity shareholder in Cullgen, Inc. and a consultant for Cullgen, Inc. J.J. was a cofounder of Cullgen, Inc., a scientific cofounder and scientific advisory board member of Onsero Therapeutics, Inc., and a consultant for EpiCypher, Inc. and Accent Therapeutics, Inc. Other authors declare no conflicts of interest.

---

**Figure S1.**
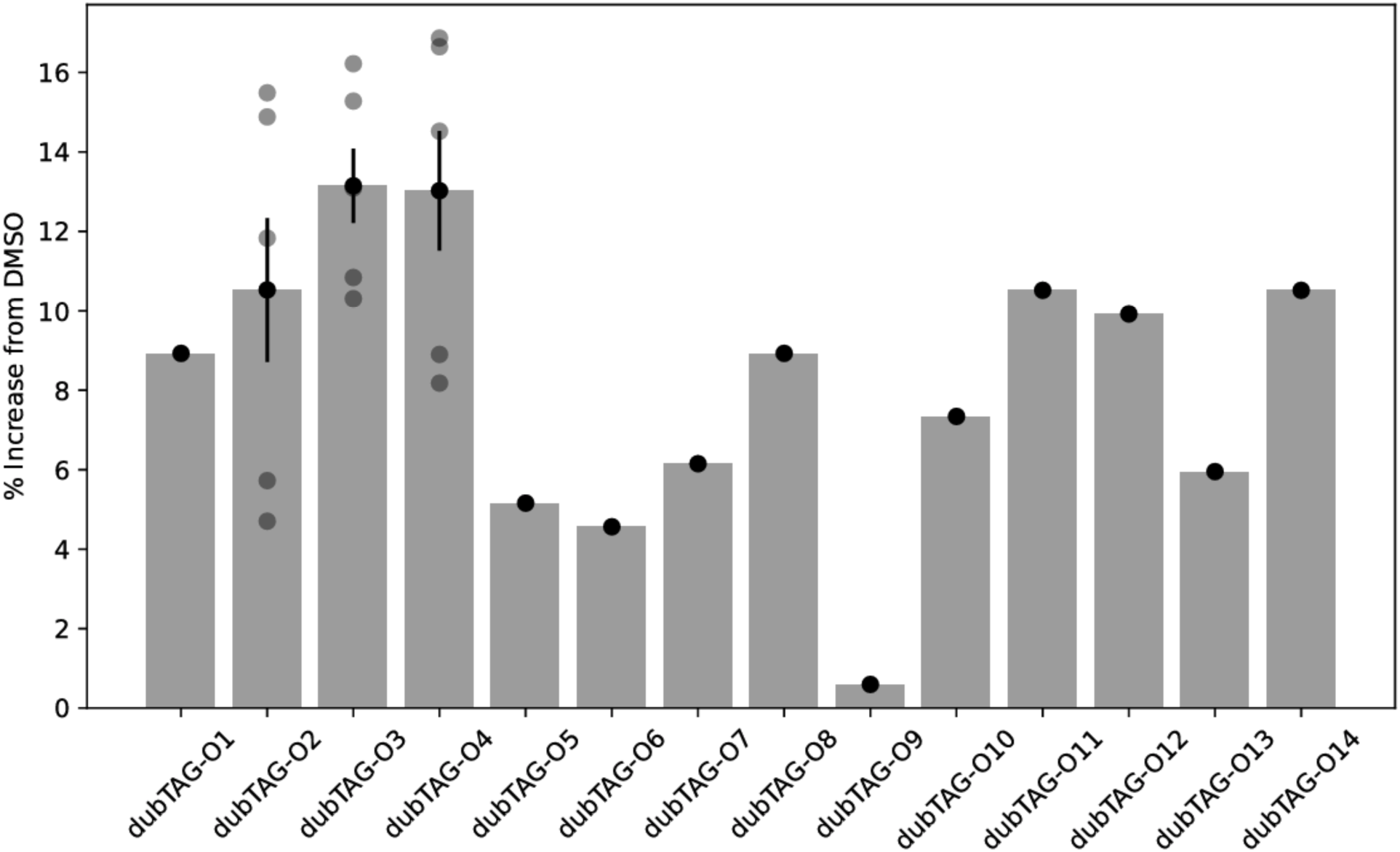
Initial screen of dubTAG effects on SOX9 levels in hESCs. Data are plotted as a percentage increase from untreated levels. All compounds were applied at 1 uM for 48h. Light points represent independent biological replicates, dark point and error bars indicate mean and standard error. Data from compounds with one biological replicate are shown for completeness and were not characterized further.

**Figure S2.**
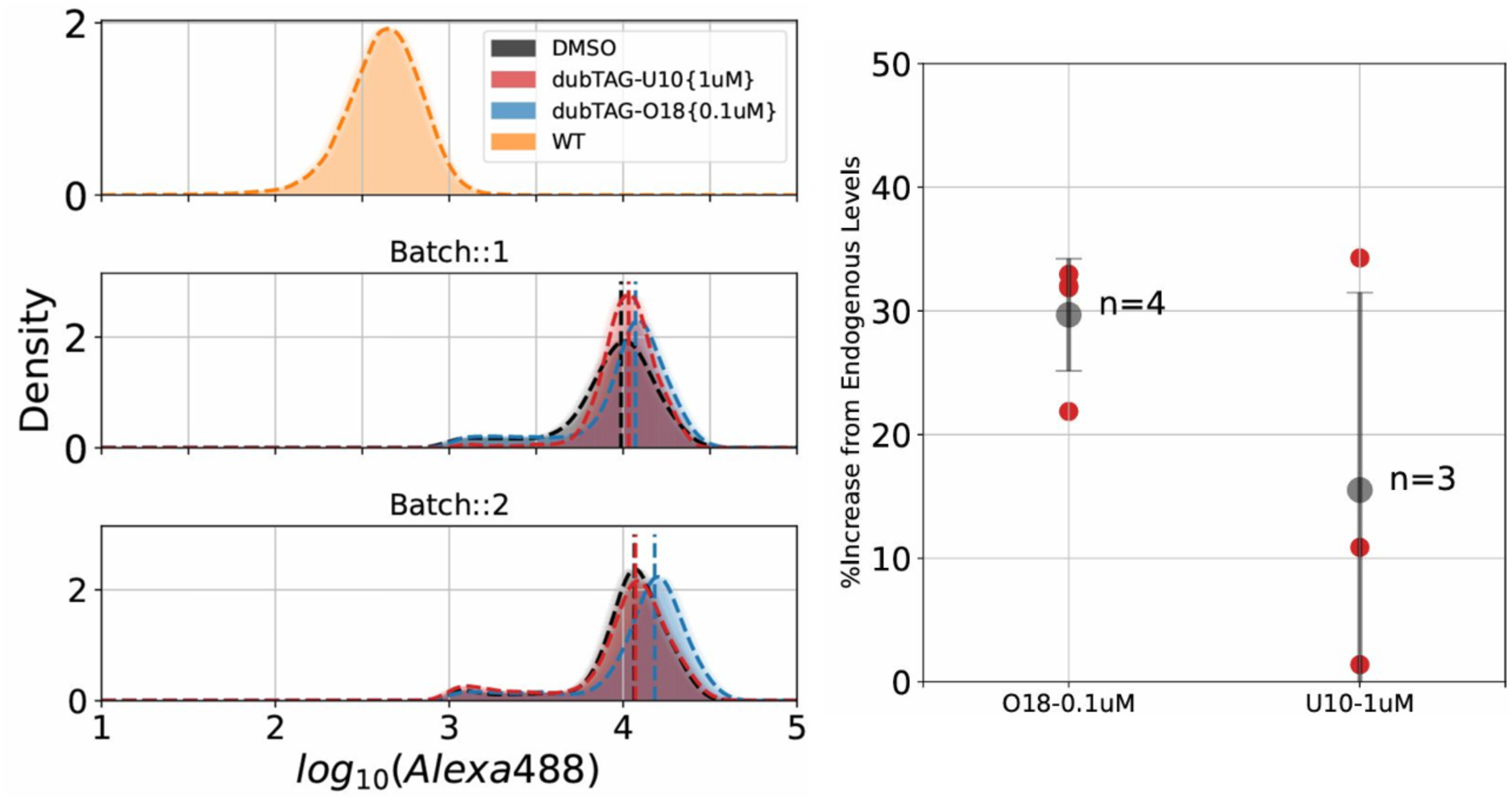
Selected dubTAGs boost endogenous levels of FKBP12^F36V^ tagged TWIST1 in CNCCs. Treatment of TWIST1-FKBP12^F36V^-V5 edited CNCCs with dubTAG-O18 and dubTAG-U10 at maximal effective concentrations as evaluated in SOX9--FKBP12^F36V^ edited CNCCs. (Left Panel) Intracellular flow-cytometry measurements across treatment conditions from two representative CNCC differentiation batches derived from TWIST1 edited h9 ESCs. (Right Panel) Quantification of TWIST1 increases from endogenous levels from dubTAG treatments. The number of replicates denote distinct differentiation batches.

**Figure S3.**
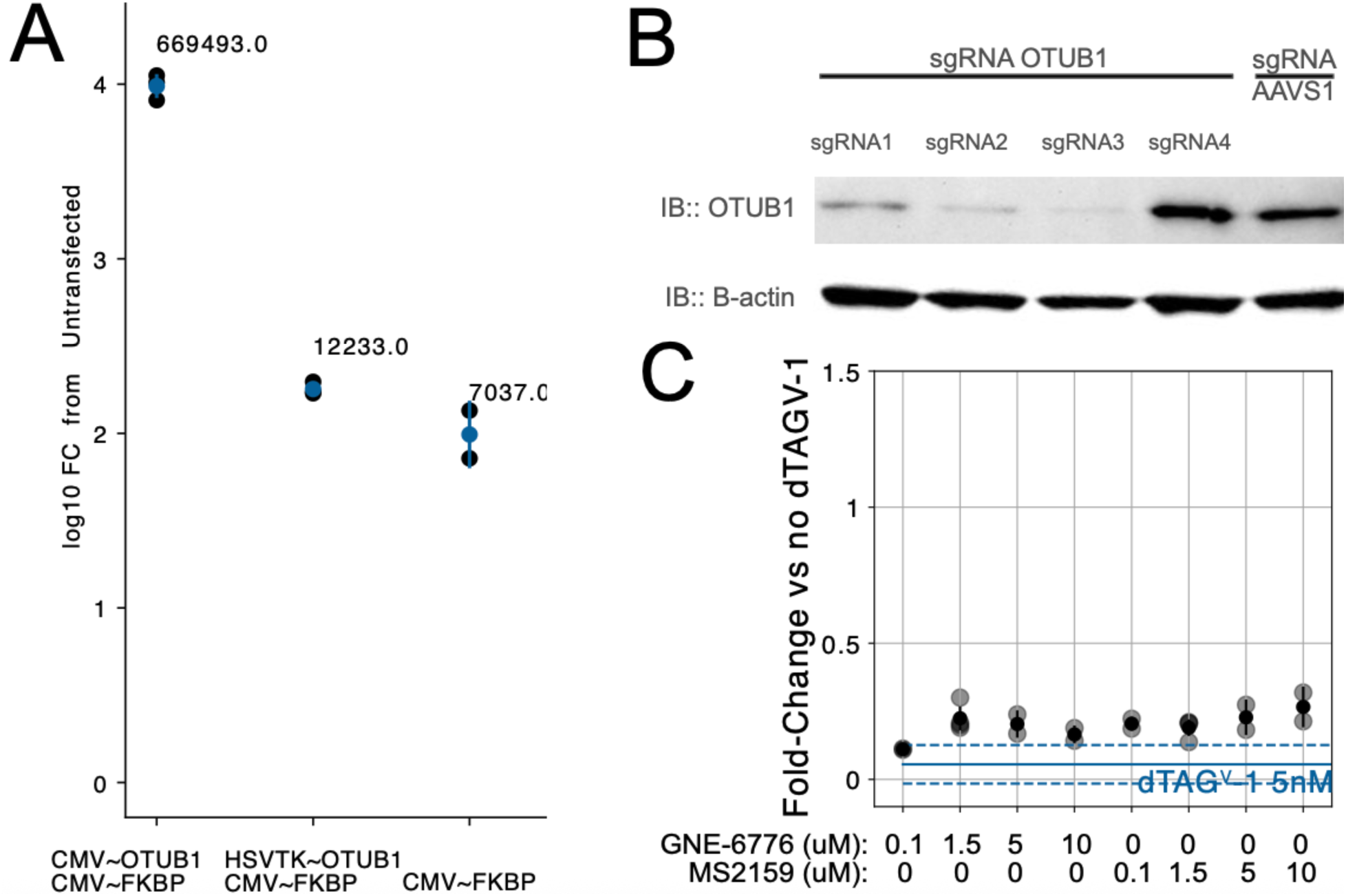
Supplementary figure related to Figure 3. (A) Change in luciferase signal (y-axis, log10 scale) relative to untransfected cells in untreated, OTUB1 WT HEK293 cells transfected with the indicated plasmids. Numbers on top of each point indicate the mean raw luciferase signal in each condition. (B) Immunoblot of HEK293 cells transfected with plasmids encoding the indicated sgRNAs targeting OTUB1 or AAVS1 (control). (C) Effect of indicated deubiquitinase binders (all 24h) on SOX9 levels following 24h 5nM dTAG^V^-1 pretreatment (of which mean and is standard error indicated by blue solid and dashed lines, respectively). Light black points represent independent biological replicates, dark points and error bars indicate mean and standard error.

**Figure S4.**
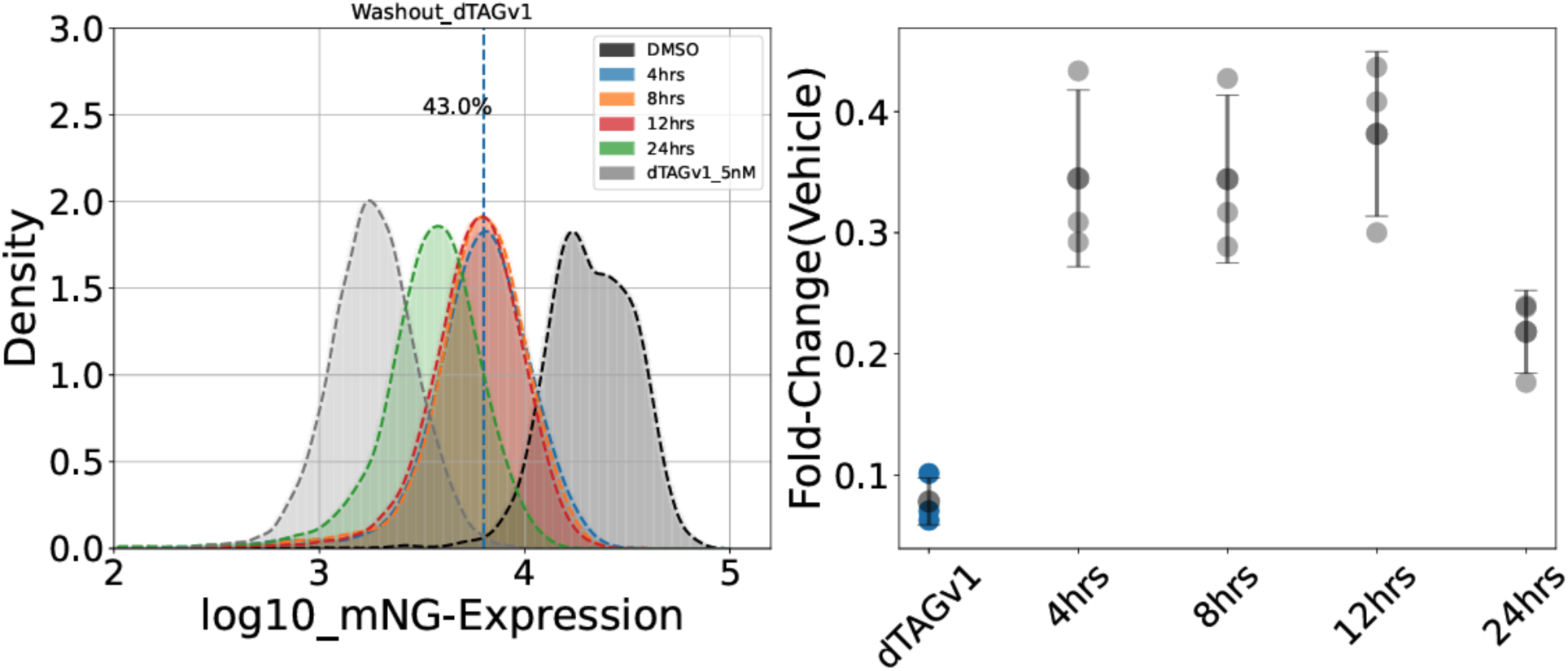
Washout of dTAG^V^-1 does not fully restore SOX9 levels. Left, representative flow cytometry histogram of mNeonGreen (SOX9) levels at distinct timepoints (colors) following washout of 5 nM dTAGV-1. Right, quantification of SOX9 level (median of flow cytometry distributions) at indicated timepoints (x-axis) following washout. Light points represent independent biological replicates, dark point and error bars indicate mean and standard error.

**Figure S5.**
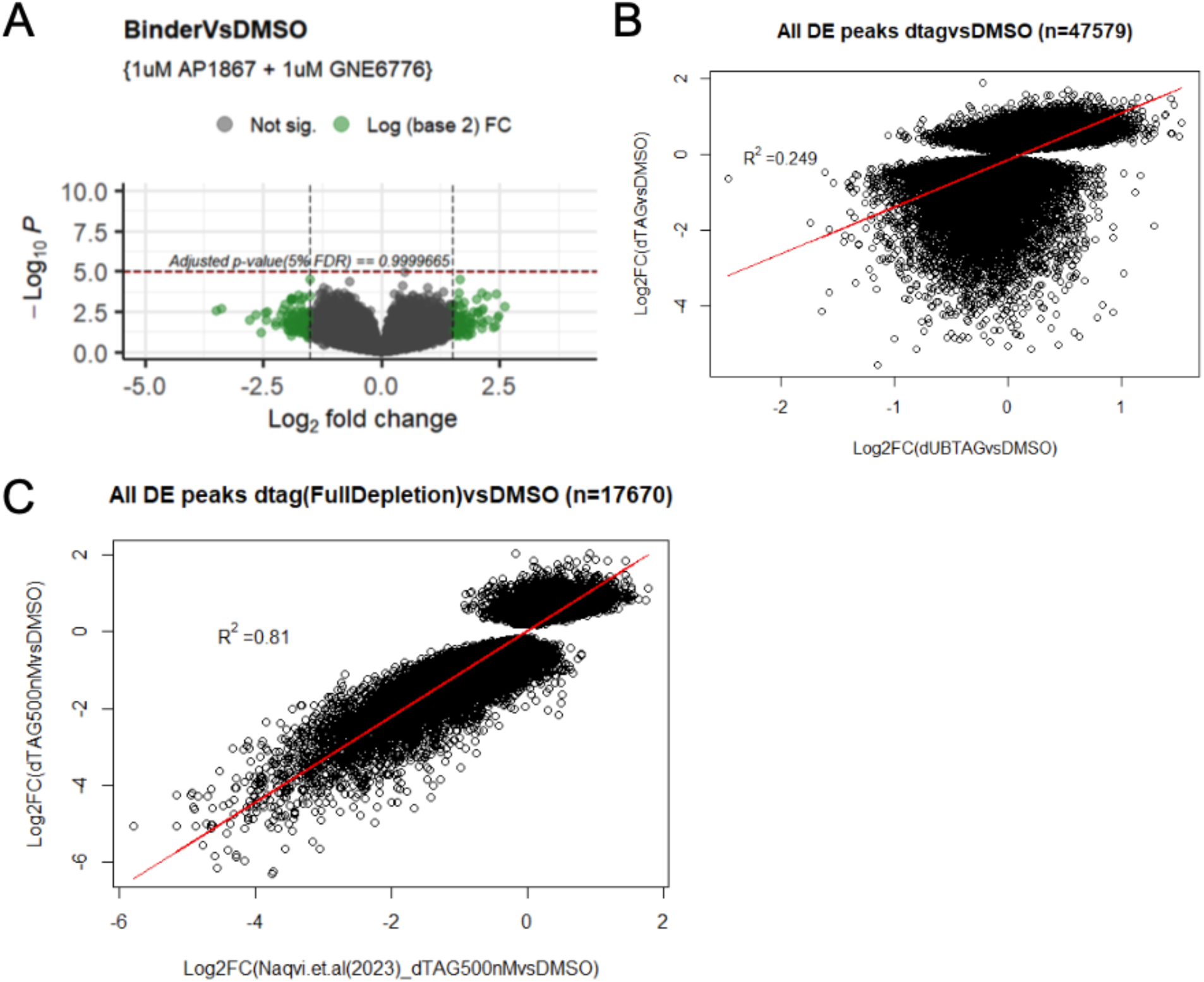
Analyses related to effects of increased SOX9 levels on chromatin accessibility in CNCCs. (A) Volcano plot of differential accessibility analysis comparing AP1867 and GNE-6776 binder treatment to DMSO. (B) Comparison of dubTAG-U10 treatment (x-axis) effect versus full SOX9 depletion effect (500 nM dTAG^V^-1) (y-axis) from this study. (C) Comparison of full SOX9 depletion effect between this study (y-axis) and Naqvi et al 2023 (x-axis).

**Figure S6.**
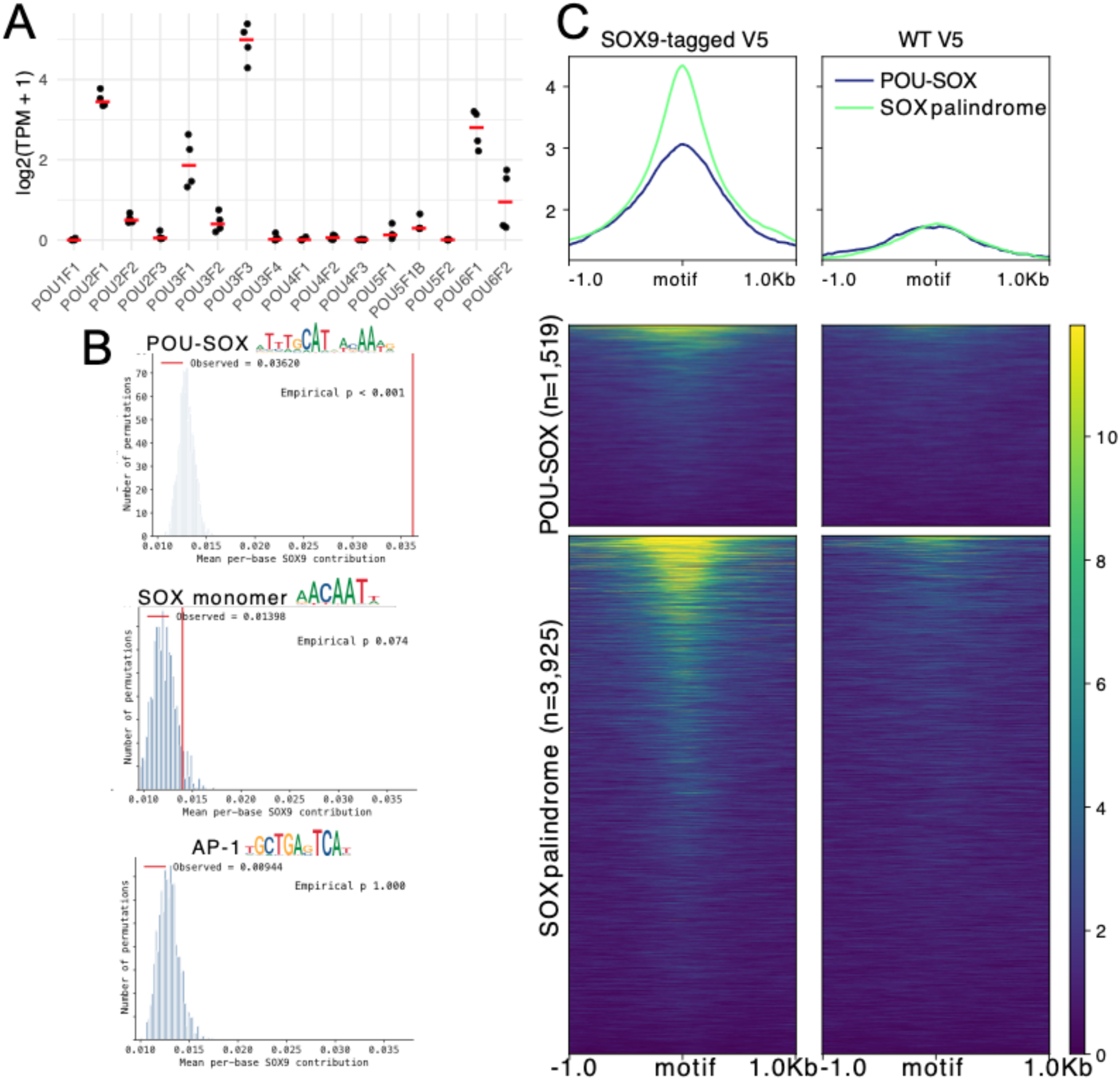
POU-SOX motif instances likely function as low-affinity SOX9 binding sites. (A) Transcripts per million (TPM) of indicated POU family TFs (x-axis) in unperturbed CNCCs (data from Naqvi et al 2025). Points represent individual biological replicates, red horizontal line represents mean. (B) Mean per-base contribution of the indicated motifs (red line), as quantified by a BPNet model trained on CNCC SOX9 (V5 ChIP-seq), compared to 1,000 permutations where motif positions were shuffled within the same REs. (C) V5 ChIP-seq signal from CNCCs with V5-tagged SOX9 present (‘SOX9-tagged V5’) or absent (‘WT’) plotted at regions centered around POU-SOX (top) or SOX9 palindrome (bottom) instances within accessible regions.

